# miR-34a negatively regulates cell cycle factor Cdt2/DTL in HPV infected Cervical Cancer Cells

**DOI:** 10.1101/2022.03.28.486030

**Authors:** Garima Singh, Sonika Kumari Sharma, Samarendra Kumar Singh

## Abstract

MicroRNAs have emerged as an important factor in regulating cell cycle and various other cellular processes. Aberration in microRNAs has been linked with development of several cancers and other diseases but still very little is known about the mechanism behind their function in regulating cellular processes. 99% of cervical cancer is caused by the infection of high-risk HPVs (HR HPVs) which suppresses multiple tumor suppressors and checkpoint factors of the host cell. It also stabilizes oncogenic proteins, one of them is Cdt2/DTL which promotes cell transformation and proliferation. In this study, we have reported that in cervical cancer cell lines miR-34a by targeting HPV E6 protein, regulates Cdt2/DTL protein level and leads to its destabilization. Destabilization of Cdt2 stabilizes onco-suppressor proteins like p21 and Set8 in these cell lines. We have also shown that the overexpression of miR-34a suppresses cell proliferation, invasion and migration capabilities of HPV positive cervical cancer cells which was reverted by overexpression of either HPV E6 or Cdt2 genes. This is the first-ever report to show that miR-34a regulates cell cycle factor Cdt2 by suppressing viral E6 protein level, which opens up the scope of exploring miR-34a as a specific therapy for cervical cancer treatment.

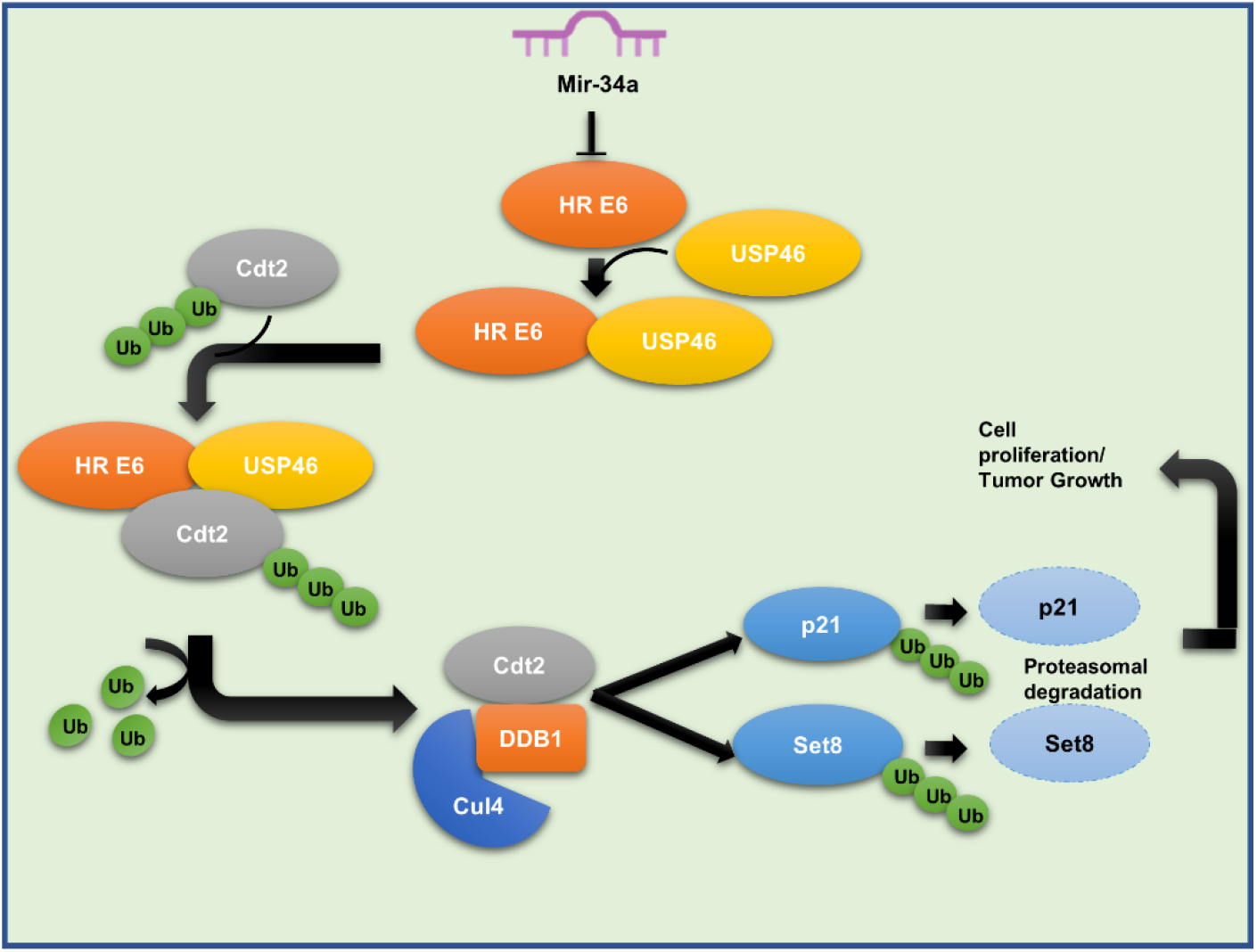

## Background

Cervical cancer is the fourth most common cause of cancer death among women with 6,04,000 new cases and 3,42,000 deaths annually (Sung *et al*, 2021). Among all the cervical cancer cases, 99% of them are attributed to human papillomavirus (HPV) infection (Moody & Laimins, 2010). Most of these cases are prevalent in developing countries like India, where due to lack of awareness, use of vaccines is negligible (no data available), therefore alternate therapeutic options could be a significant solution. Apart from cervical cancer, HPV is also known to cause penile, vulvar, anal, vaginal, oropharyngeal and head and neck cancers (Moscicki *et al*, 2012).

HPV is a non-enveloped double-stranded DNA virus which eventually integrates into the host genome upon infection. It infects the basal cutaneous squamous epithelium and causes oncogenic transformation and hyperproliferation of the cells upon genomic integration (zur Hausen, 2002). Till date, 15 HPVs have been categorized as the high risk (HR) out of which HPV 16 and HPV 18 are responsible for almost 70% of cervical cancer cases (Moody & Laimins, 2010). The HR HPV encoded oncoproteins E5, E6 and E7 are the primary factors responsible for oncogenic transformation and progression of cervical cancer. The HR E7 interacts with retinoblastoma family proteins (RB1, RBL1 and RBL2) and degrades them to release E2F (Boyer *et al*, 1996; Münger *et al*, 2001) whereas HR E6 causes ubiquitin-mediated degradation of p53 via E6AP, ubiquitin ligase (Scheffner *et al*, 1990). Apart from regulating multiple genes and proteins, HR E6 also inhibits apoptotic signaling and activates telomerase reverse transcriptase to promote immortalization of cells (Moody & Laimins, 2010).

Cdt2/DTL (denticleless) is an essential mammalian protein which acts as a substrate adaptor for a cullin RING E3 ligase system, CRL_4_. CRL_4Cdt2_ is recognised as one of the master regulators of cell cycle progression and genome stability (Abbas & Dutta, 2011). It ensures the timely degradation of multiple cell cycle factors like p21, Set 8 and Cdt1 etc. which are involved in DNA replication initiation, progression, apoptotic checkpoint regulation and chromatin modification (Jin *et al*, 2006; Hyun *et al*, 2010; Jørgensen *et al*, 2011), in a replication-coupled and PCNA-dependent manner during S phase or after DNA damage (Dar *et al*, 2014). The Cdt2 protein level is maintained in the cell by factors like CRL1^FBXO11^, CRL4^DDB2^, APC/C-Cdh1 system etc., it increases during G1 to S transition and then decreases during mitosis (Wu *et al*, 2021). Cdt2 is highly amplified in various cancers like lung cancer, breast cancer, colon cancer, Erwig sarcoma, cervical cancer etc. (Wu *et al*, 2021). In case of HR HPV mediated cervical cancers E6 protects Cdt2 from ubiquitin-mediated degradation by recruiting a deubiquitinase, USP46 to deubiquitinate and stabilise Cdt2 (Kiran *et al*, 2018).

MiRNAs (micro RNAs) are small non-coding regulatory RNAs of size 17-25 nucleotide in length which modulates the expression of more than 60% of genes at the post-transcriptional level in mammals (Friedman *et al*, 2009). MiRNA either degrades mRNA or inhibits translation by predominantly targeting 3’UTR (Untranslated Region) sequences of mRNA complementary to its functional 8-base seed sequence (Sotiropoulou *et al*, 2009). Thus, miRNA can be either tumor promoter or suppressor based on its target mRNA. In human genome, ∼416 miRNA genes encode for ∼340 distinct mature miRNAs known to date. However, the molecular targets and the mechanism behind their functions for most of the miRNAs are yet to be discovered. Mutation or deletion in miRNA genes and their aberrant expression have been implicated in many diseases including cancers but the mechanism behind is poorly understood. Among all these miRNAs, miR-34a which is located on chromosome 1p36 is believed to be a tumor suppressor and has been reported to be downregulated in several cancers including cervical cancer (Lodygin *et al*, 2008; Wang *et al*, 2008; Li *et al*, 2014; Rokavec *et al*, 2014; Si *et al*, 2019). Earlier it has also been shown that p53 positively regulates miR-34a (Tarasov *et al*, 2007). However, the pathological and clinical relevance of miR-34a in cervical cancer is still not well understood. In this study, we have explored the role of miR-34a in cervical cancer cell lines. We have found that Cdt2 is the novel target of miR-34a in HPV positive cervical cancer cells. It regulates Cdt2 expression by targeting E6 which is responsible for stabilization of Cdt2 in HR HPV infected cervical cancer. miR-34a expression selectively affects proliferation in these cancer cells while not much effect is seen in non-cancerous cell lines. This is the first report, To the best of our knowledge, of miR-34a destabilizing Cdt2 by inversely regulating E6. Hence, this study could pave way for future research in exploring the role of miR-34a as an alternate therapy for cervical cancer.

## Materials and Methods

### Cell Lines and Media

Cervical cancer cell lines (HPV positive) SiHa and HeLa, purchased from the American Type Culture Collection (ATCC, USA) and cervical cancer cell line (HPV negative) C33A and human kidney cell line HEK293T, purchased from the National Centre for Cell Science (NCCS, Pune, India) were grown in Dulbecco’s Eagle Modified Medium (DMEM, Gibco, USA) supplemented with 10% Fetal Bovine Serum (FBS, Gibco, Brazil) and 1% Pen-Strep (Penicillin-Streptomycin, Gibco, USA). The cells were cultured at 37°C, with 95% humidity and 5% CO_2_.

### Plasmids and miRNA

Empty plasmid backbone pcDNA 3.1 was acquired from Addgene, Watertown, USA and miR-34a plasmid was gifted by Moshe Oren, The Weizmann Institute of Science, Israel. Flag-Cdt2 and 16E6 plasmids were gifted by Dutta’s Lab, University of Virginia, USA. Other microRNA mimics were acquired from Dharmacon, USA.

### Cell Transfection

HEK293T, C33A, HeLa and SiHa cells were seeded (1×10^5^) in 6 well plates 24 hours prior to the transfection. The plasmid vectors (miR-34a, Flag-Cdt2 and 16E6) were transfected to the cells with turbofect™ (Thermo Fisher Scientific, Massachusetts, USA) the following day according to the manufacturer’s protocol. The cells were cultured at 37°C, with 95% humidity and 5% CO_2_ for 48 hours.

### Western Blot Analysis

Cells were harvested and total protein was extracted using Radioimmunoprecipitation assay (RIPA) buffer (50 mM Tris-Cl pH 7.5, 150 mM NaCl, 1% Nonidet P-40, 0.5% sodium deoxycholate,.1% sodium dodecyl sulfate, 1 mM PMSF) supplemented with protease inhibitor cocktail (Cell Signaling Technology, Massachusetts, USA). The protein concentration was measured using Bradford protein assay (Sigma, Missouri, USA). The total protein extract was separated via SDS-PAGE and then transferred onto the 0.22 µm or 0.45 µm PVDF membrane (Millipore, Massachusetts, USA). The membrane was blocked with either 5% BSA (SRL, Mumbai, India) or 5% fat-free milk in TBST (Amersham, GE, Chicago, USA) for 1 hour at 4°C followed by 1 hour at room temperature with gentle shaking. The membrane was then washed and probed with primary antibody overnight as per manufacturers’ instructions followed by washing and then incubating in secondary antibody (in TBST with 2.5% fat-free milk). The blot was then developed by enhanced chemiluminescence (ECL) substrate (Bio-Rad, Callifornia, USA) and documented using the chemidoc (Bio-Rad XRS+).

### Cell Migration and Cell Invasion Assays

Post 48 hours of transfection, cells were washed with PBS and resuspended in serum-free DMEM at 5×10^5^ cells/ml. For invasion assay, 1×10^5^ cells were placed in upper chamber of transwell plate (Corning, New York, USA), precoated with 2 mg/ml ECM gel (Sigma-Aldrich, Missouri, USA) and DMEM supplemented with 10% fetal bovine serum was added to the lower chamber. Cells were incubated in CO_2_ incubator for 48 hours. The remaining cells from upper chamber were removed and the cells invaded through the membrane were fixed with 5% glutaraldehyde. Invaded cells were stained with 0.2% crystal violet solution in 2% ethanol and counted under inverted microscope.

For migration assay, 1×10^5^ cells were placed in upper chamber of the transwell plate and DMEM supplemented with 0.5% FBS and 40ug/ml collagen I (Sigma-Aldrich, Missouri, USA) was added to the lower chamber. Cells were incubated for 24 hours in CO_2_ incubator. The remaining cells were removed from upper chamber and the migrated cells were fixed with 5% glutaraldehyde. The migrated cells were stained with 0.2% crystal violet solution in 2% ethanol and counted under inverted microscope.

### Immunofluorescence

Post 48 hours of transfection, 1×10^4^ cells were plated on glass coverslips placed in 6 well plate and incubated with DMEM medium containing 10% FBS for 24 hours in CO_2_ incubator. Cells were then washed with PBS and fixed with 4% paraformaldehyde in PBS for 20 mins. Fixed cells were permeabilized with 0.1% Triton X-100 in PBST and blocked with 5% BSA for 1 hour. Coverslips were incubated with primary antibody overnight at 4°C in dark followed by incubation with secondary antibody at 4°C in dark humid chamber for 1 hour. The coverslip was mounted on slide using mounting media (Vectashield, Vector Laboratories, San Francisco) and observed under confocal microscope (STED, Leica, Wetzlar, Germany).

### Statistical Analysis

All the western experiments were performed in biological triplicates. The immunofluorescence, growth curve, migration and invasion assays were performed in experimental triplicates. All the data are presented as the mean ± SD and an unpaired student’s t-test was performed to calculate the significant value. Image J software was used to quantify the intensity of protein bands wherever applicable.

## Results

### miR-34a negatively regulates Cdt2 protein stabilization in cervical cancer cells

Previous studies have shown that in HR HPV infected cervical cancer cells, Cdt2 is stabilized by HPV E6 protein (Kiran *et al*, 2018). Also, it has been well established that miR-34a is downregulated in cervical cancer (Wang *et al*, 2008). In order to investigate whether there is a correlation between miR-34a and expression of Cdt2, we transfected HEK293T cells (non-cancerous cell line) and two HR HPV positive cervical cancer cell lines SiHa and HeLa with miR-34a. A significant reduction in expression of Cdt2 was detected in both the cervical cancer cell lines (Fig.1A Lane 3,4,5 and 6 and Fig.1B), whereas ectopic expression of miR-34a did not cause any significant change in expression of Cdt2 in HEK293T cell lines (Fig.1A Lane 1and 2 and Fig.1B).

**Fig. 1:**
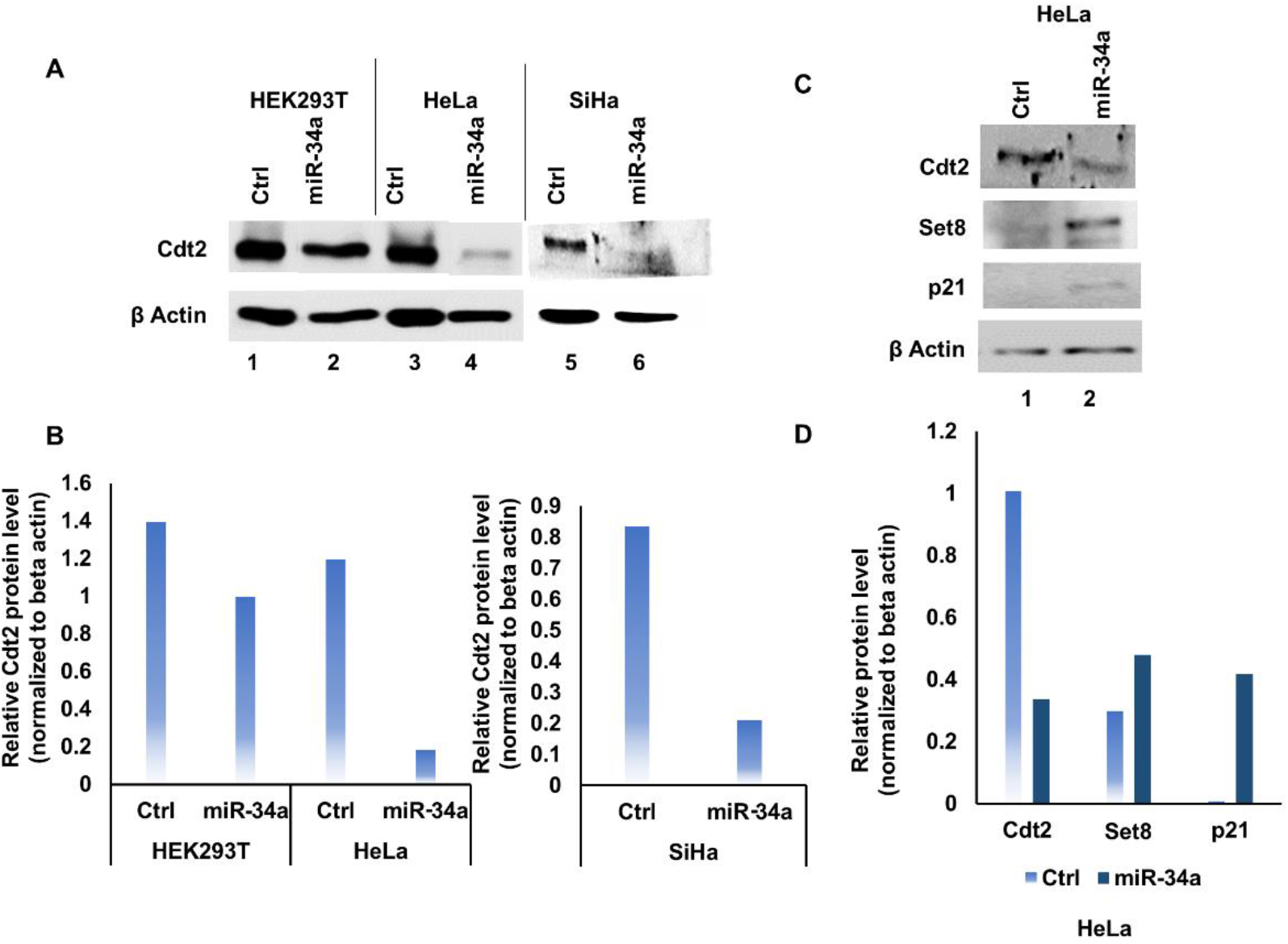
**Overexpression of miR-34a decreases Cdt2 expression level in cervical cancer cells. A. Western blot analysis was performed for Cdt2 protein level upon 48h treatment with miR-34a in HPV positive cells (HeLa and SiHa) and HEK293T cells. B. Quantification of the Cdt2 protein level in HEK293T, HeLa and SiHa cell lines upon transfection with miR-34a for 48h in comparison to their respective vector control. C. Western blot analysis was performed for p21 and Set8 protein levels upon 48h treatment with miR-34a in HPV positive cells (HeLa cells) in comparison to the cells transfected with control. D. Quantification of SET 8 and p21 protein level in HPV positive cell line upon transfection with miR-34a for 48h in comparison to vector control.**

Since Cdt2 facilitates degradation of p21 and Set 8 (Abbas & Dutta, 2011; Kiran *et al*, 2018), we next checked the level of p21 and Set 8 upon miR-34a expression in HR HPV positive cervical cancer cells and found that Cdt2 destabilization stabilizes both p21 and Set 8 protein level (Fig. 1C and 1D).

### miR-34a decreases Cdt2 expression in cervical cancer cells by targeting HPV protein E6

We have shown that miR-34a destabilizes Cdt2 level in HPV positive cervical cancer cells but does miR-34a also destabilizes Cdt2 in HPV negative cervical cancer cells too? To answer this question, we transfected C33A (non-HPV cervical cancer) cells with miR-34a expressing plasmid. Surprisingly we discovered that miR-34a expression does not suppress Cdt2 level in the C33A cell line (rather it increases Cdt2 level slightly, Fig. 2A and 2B).

**Fig. 2:**
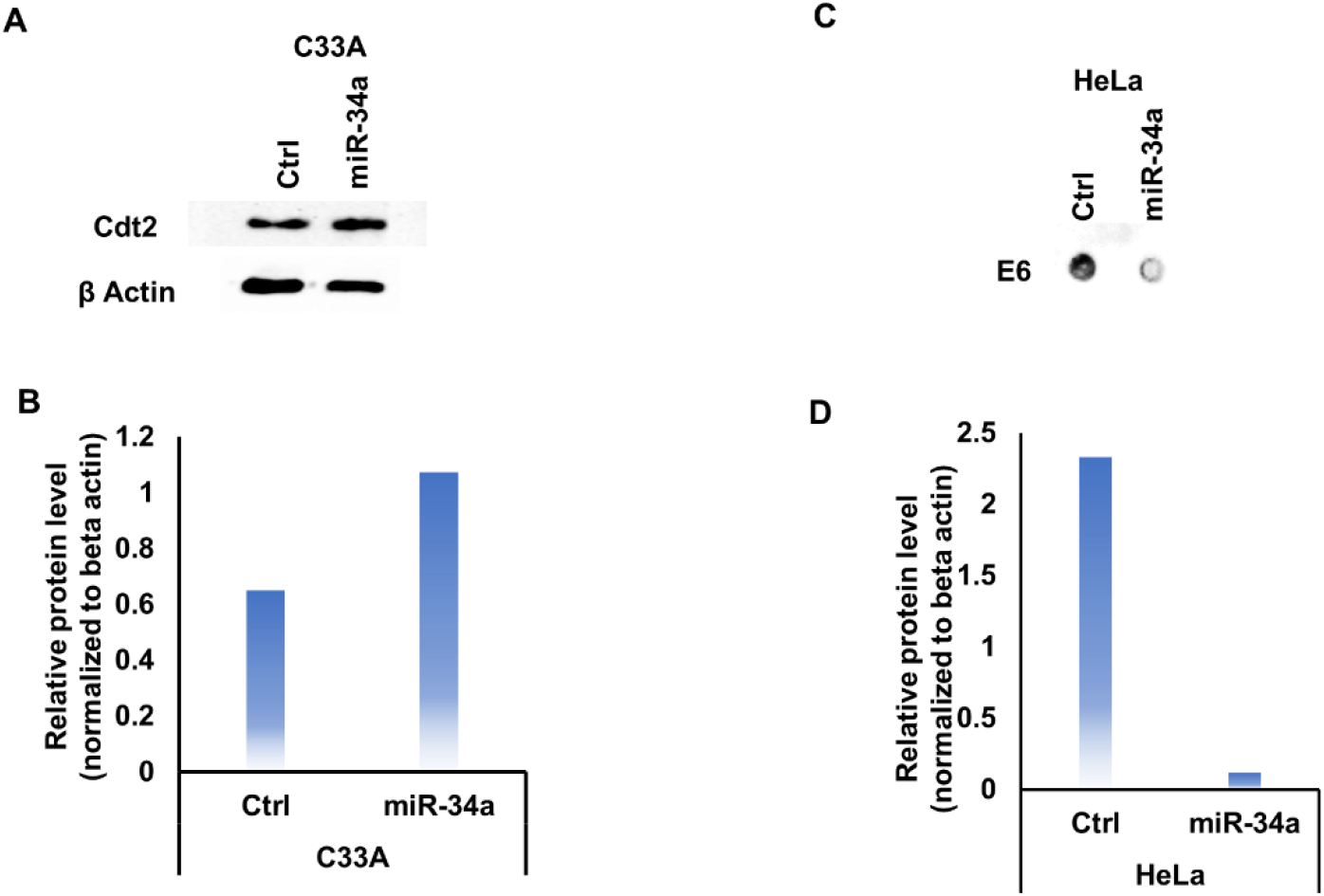
**MiR-34a decreases Cdt2 expression in HPV infected cervical cancer cells by targeting HPV E6. A. Western blot analysis was performed for Cdt2 protein level upon 48h treatment with miR-34a in HPV negative cervical cancer cells (C33A). B. Quantification of the Cdt2 protein level in C33A cell line upon transfection with miR-34a for 48h in comparison to vector control. C. Dot blot analysis was performed for E6 protein level upon 48 h after transfection in HPV positive cell (HeLa) with miR-34a in comparison to relative control. D. Quantification of E6 protein level in HPV positive cell line (HeLa) upon transfection with miR-34a for 48h in comparison to vector control.**

Since E6 is the major regulator in HPV infected cells, we wanted to investigate the effect of miR-34a on E6 expression. The dot blot profile of E6 protein in the HeLa cells shows that miR-34a expression reduces the E6 protein level (Fig.2C and 2D), suggesting that miR-34a/Cdt2 axis is directly regulated by E6 suppression in cervical cancer.

### miR-34a suppresses the proliferation of HPV infected cervical cancer cells

We have already shown that miR-34a destabilizes Cdt2 and E6 which are one of the major players of proliferation and tumor growth in HR HPV infected cervical cancer cell lines. Therefore, we wanted to see what role miR-34a overexpression plays in growth and proliferation of cervical cancer cells. We observed that mir-34a expression significantly inhibits the growth of cervical cancer cell lines (Fig. 3A and 3B). While in case of HEK293T cells, miR-34a overexpression decreases the cell proliferation initially (at 48 hours) which get normalized by 72 hours (Fig. 3C).

**Fig. 3:**
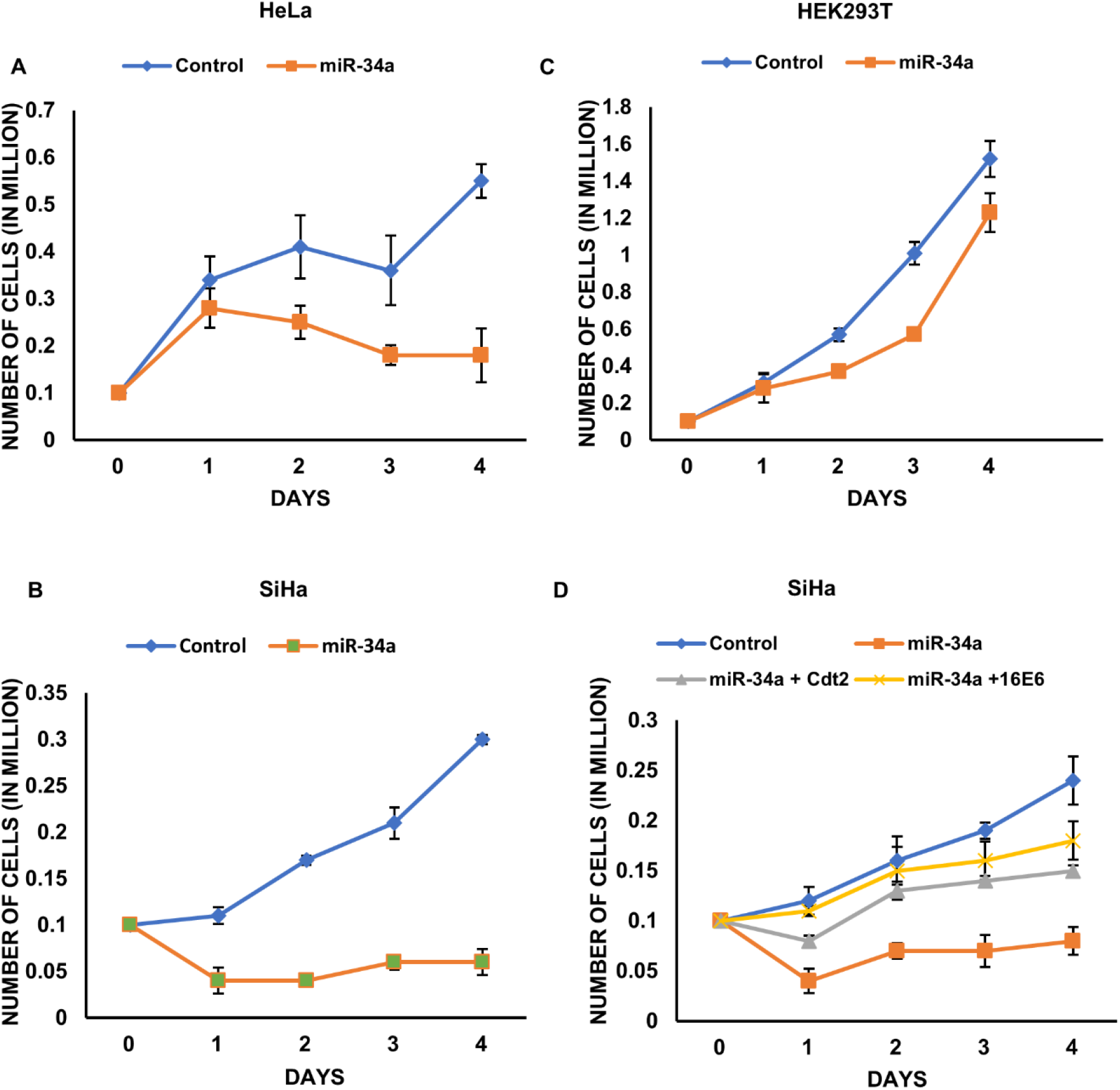
**Ectopic expression of miR-34a suppresses cell proliferation in HPV positive cervical cancer cells. A. Growth curve for HeLa-miR-34a shows the reduction in growth in comparison to relative control. B. Growth curve for SiHa-miR-34a shows the reduction in growth in comparison to relative control. C. Growth curve for HEK293T-miR-34a shows the growth inhibition which is compensated by day 4. D. 0.1 million SiHa cells were seeded 24 hours prior to transfection with mock, miR-34a, miR-34a+Cdt2 and miR-34a+16E6 and cell growth were observed every 24 hours for 4 days. Each value represents the mean of three readings. Error bar represents S.D.**

Further, we wanted to see if decreased proliferation rate of HPV positive cells upon miR34a expression could be normalized by overexpression of Cdt2 or E6 proteins. For this, we transfected SiHa cells with either miR-34a alone or miR-34a in combination with 16E6 plasmid or miR-34a in combination with Flag Cdt2 plasmid. The results showed that ectopic expression of 16E6 or Cdt2 could rescue the growth of cervical cancer cells (Fig. 3D). However, the rescuing of inhibited proliferation is more in case of miR-34a and 16E6 co-transfection compared to miR-34a and Flag-Cdt2 co-transfection. We had also co-transfected SiHa cells with all the three vectors containing miR-34a, Flag-Cdt2 and 16E6 but it proved to be toxic and could not rescue the growth inhibition caused by miR-34a expression (Fig. S1).

### miR-34a expression causes morphology change and cell growth suppression

To observe the morphological changes occurring in the cervical cancer cells upon miR-34a transfection we performed phase-contrast microscopy and observed that the untreated HeLa and SiHa cells maintained their morphology and were in close contact with each other even after 48 hours of incubation. In contrast, the miR-34a treated cells lost their original shape after 48 hours of treatment (Fig. 4A). Also, miR-34a transfected HeLa and SiHa cells have lower confluency than the control (Fig. 4A), suggesting decreased proliferation rate.

**Fig. 4.**
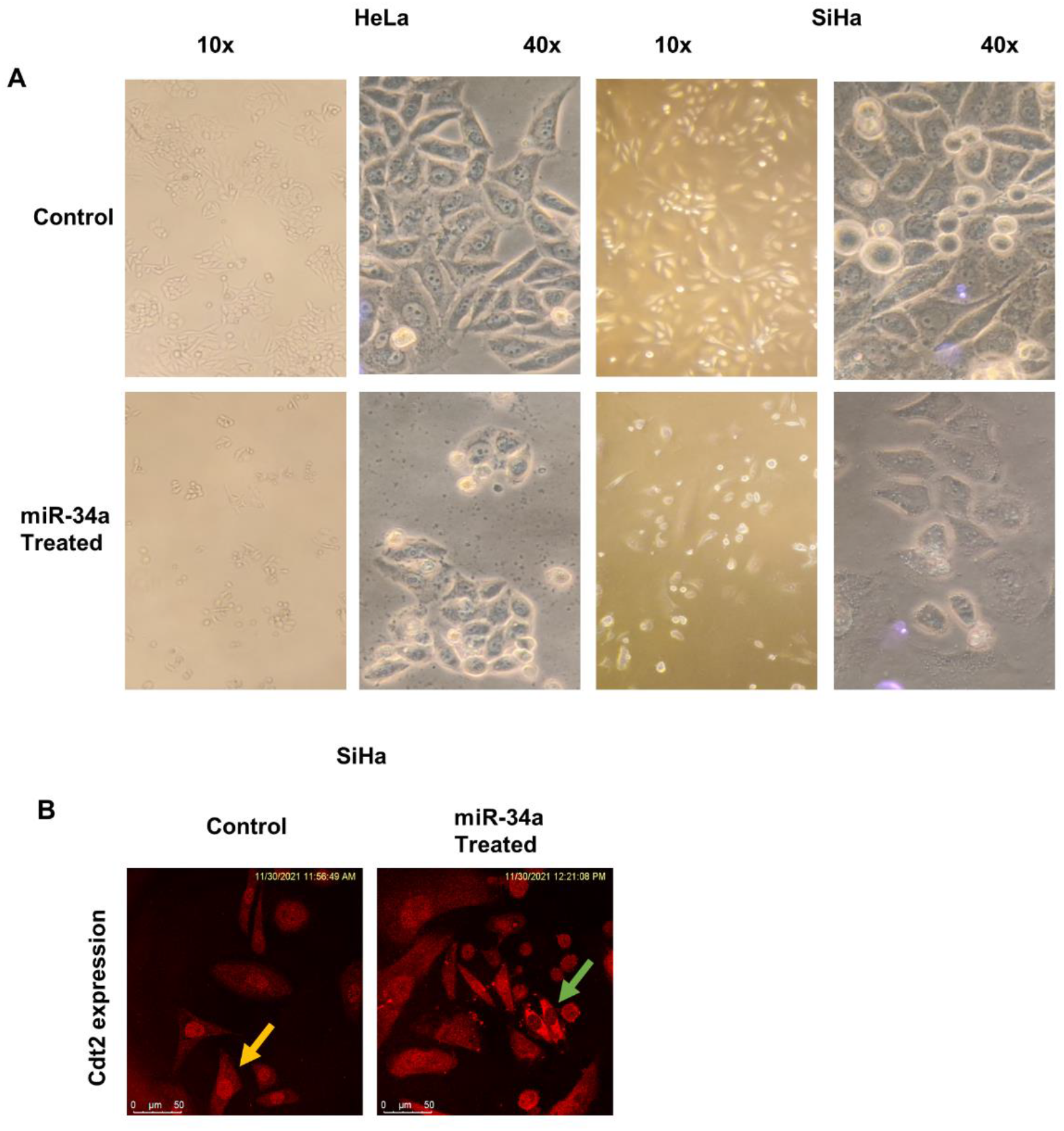
**MiR-34a expression causes morphology change. A. HeLa and SiHa cells were transfected with miR-34a for 48 hours and phase-contrast microscopy was performed at 10X and 40X magnifications. B. SiHa cells were transfected with miR-34a for 48 hours, probed with anti-Cdt2 proteins and confocal microscopy was performed. Scale bar, 50 µm.**

Cdt2 is involved in G1 to S phase transition of the cell cycle and is majorly localized in the nucleus (Huh & Piwnica-Worms, 2013). By confocal microscopy, it was evident that Cdt2 is enriched in the nucleus of control SiHa cells but get exported to the cytoplasm upon miR-34a overexpression (green arrow vs yellow arrow Fig. 4B). Also, the miR-34a treated HPV positive cervical cancer cells showed a higher number of dead cells.

### Upregulation of miR-34a suppresses both cell migration and invasion of HPV infected cervical cancer cell lines

In order to determine what role miR-34a plays in cell proliferation, metastasis and invasion of HPV positive cervical cancer cells, transwell invasion and migration assays were performed. The invasive potential of HeLa and SiHa cells transfected with miR-34a gets reduced by 24.6% and 45.3% respectively, in comparison to control transfected cells (Fig.5A-5D). we also observed that the migration level of HeLa cells was decreased by ∼ 0.5-fold after transfection with miR-34a, compared to control (Fig.5E and 5F). Hence, both invasion and migration were significantly suppressed by overexpression of miR-34a.

**Fig. 5:**
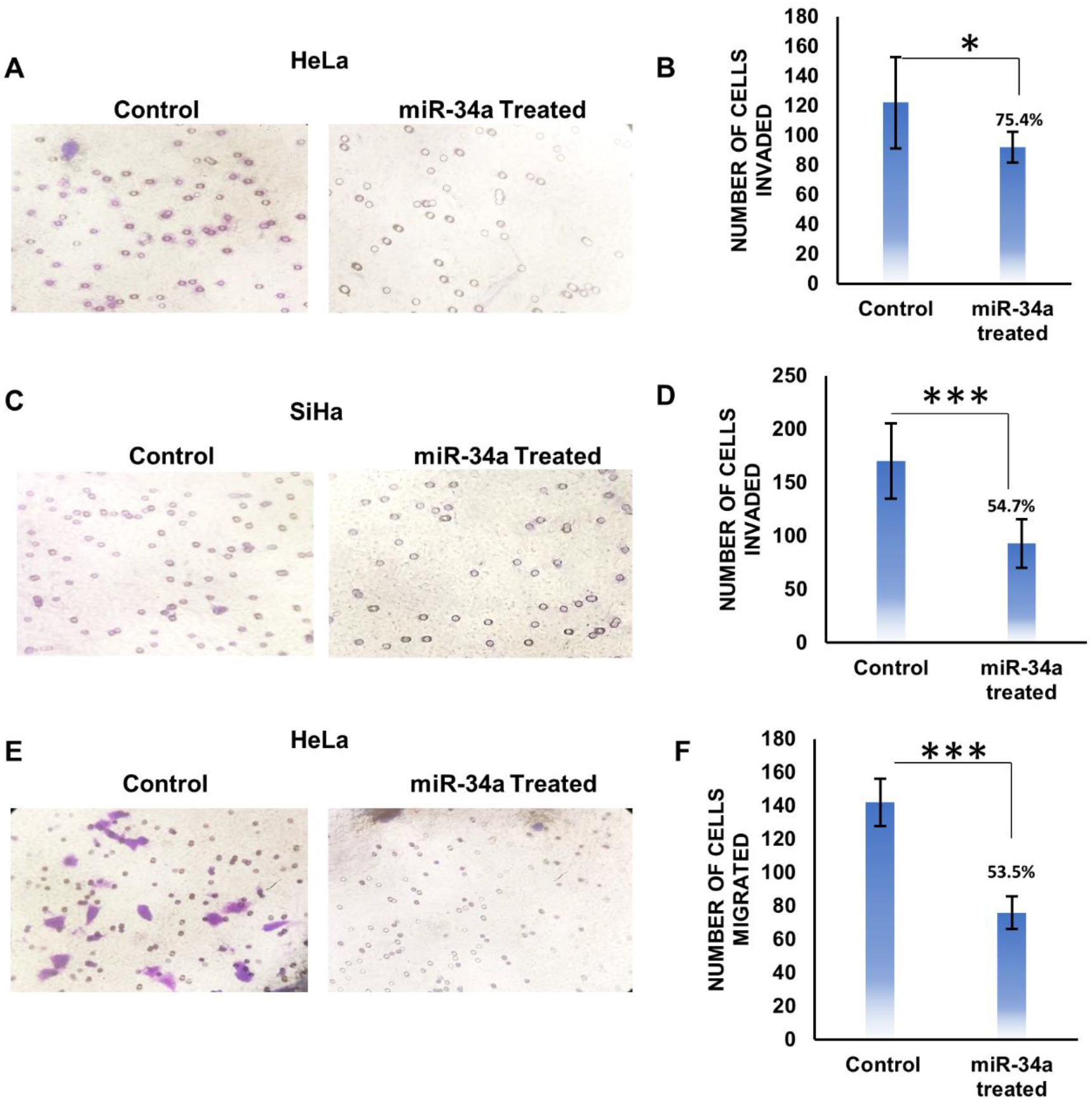
**Effect of miR-34a expression on transwell migration and invasion. A. Representative images of transwell invasion assay of HeLa cells with ectopic expression of miR-34a and respective vector control. B. Graphical representation of transwell invasion assay of HeLa cells with ectopic expression of miR-34a (92±10.38), showing the reduction in transwell invasion in comparison to vector control (122±30.64). C. Representative images of transwell invasion assay of SiHa cells with ectopic expression of miR-34a and respective vector control. D. Graphical representation of transwell invasion assay of SiHa cells with ectopic expression of miR-34a (93±22.74), showing the reduction in transwell invasion in comparison to vector control (170±35.40). E. Representative images of transwell migration assay of HeLa cells with ectopic expression of miR-34a and respective vector control. F. Graphical representation of transwell migration assay of HeLa cells with ectopic expression of miR-34a (76±9.66), showing the reduction in transwell migration in comparison to vector control (142±14.29). 8 different areas were selected randomly and number of invaded and migrated cells were counted. Error bar represents S.D. * represents p value < 0.05, *** represents p value < 0.001.**

## Discussion and Conclusion

Dysregulation of miRNAs has been linked with various cancers (Garzon *et al*, 2009; Feng *et al*, 2015); these biomolecules can act either as tumor suppressors or tumor promoters (Okada *et al*, 2014; Sachdeva *et al*, 2014). Cervical cancer which is one of the deadliest cancers in women worldwide (fourth-largest killer among cancers) is caused by high-risk HPV infection which in turn interferes with many cellular processes and drives the infected cells towards oncogenic transformation. Several miRNAs have been reported to be misregulated in cervical cancer cells (both in patient samples and cell lines data; Gocze *et al*, 2013; Honegger *et al*, 2015; Veena *et al*, 2020) and one of them is miR-34a, which is encoded on chromosome 1p36 (Li *et al*, 2014; Rokavec *et al*, 2014) in humans. miR-34a has been shown to play a critical role in maintaining the healthy functioning of cells by acting as a tumor suppressor in many solid tumors. It has been reported to repress over 700 transcripts involved in cellular proliferation, survival and plasticity (Slabáková *et al*, 2017), however, very little is known about the mechanisms involved in this process.

In this study, we have discussed role of miR-34a in regulation of a master cell cycle factor Cdt2/DTL which is an essential protein involved in G1 to S phase transition (Abbas & Dutta, 2011; Rizzardi & Cook, 2012; Barr *et al*, 2017). We have shown that miR-34a decreases the level of Cdt2 in HPV positive cervical cancer cells (Fig. 1) by suppressing viral E6 protein (Fig 2C). This is the first-ever study to show that HR HPV E6 oncoprotein, which suppresses many tumor inhibitor proteins, is directly regulated by miR-34a.

HR E6 oncoprotein hijacks many cell cycle pathways and prevents the host cell from going in cell cycle arrest and apoptosis in response to the genotoxic viral infection (Moody & Laimins, 2010; Kiran *et al*, 2018). It stabilizes Cdt2 in HR HPV cell lines, which in turn proves to be essential for survival of cancerous cells both in-vitro and in-vivo (Kiran *et al*, 2018). Cdt2 is an adaptor protein of CRL4 E3 ubiquitin ligase complex and is involved in proteasomal degradation of cell cycle factors like p21, Cdt1, Set8 etc. for the progression of G1/S and S phase to the M phase (Abbas & Dutta, 2011). Silencing of E6 gene (by siRNA) has been shown to promote polyubiquitination of Cdt2 which eventually leads to degradation of the same (Kiran *et al*, 2018).

In this study we observe destabilization of Cdt2 by miR-34a is specific to HPV positive cervical cancer cell lines (Fig 1A Lane 3,4,5,6 and 1B) while no change is observed in non-cervical cancer cell line (Fig. 1A lane 1,2 and 1B) or HPV negative cervical cancer cell line (Fig. 2A and 2B), which indicates that Cdt2 is not a direct target of miR-34a. Since Cdt2 is a major component of CRL4-E3 Ubiquitin ligase system, therefore, destabilization of Cdt2 leads to stabilization of p21 and Set8 which otherwise is present in very low levels in HPV positive cervical cancer cells (Fig.1D and 1E). In earlier studies, ectopic expression of miR-34a has been shown to arrest the cell cycle at G1/S and G2/M phase in cervical cancer cells (Wang *et al*, 2009). Our results explain the above finding that it might be due to elevated levels of p21 and Set8 which are pro-apoptotic and anti-cancerous proteins (Houston *et al*, 2008; Abbas & Dutta, 2011).

Further, for the first time to our knowledge we have shown that miR-34a overexpression decreases the HPV E6 protein expression (Fig. 2B and 2C) which stabilizes Cdt2 by importing a deubiquitinase USP46 from cytoplasm to nucleus (Kiran *et al*, 2018). Suppression of E6 by miR-34a provides favorable opportunity for CRL1^FBXO11^ or CRL4^DDB2^ E3 ubiquitination mediated degradation of Cdt2. Although it has been well established that miRNA can degrade or inhibit the target mRNA by interacting with the 3’UTR of the target sequence (Fang *et al*, 2015) in mammalian cells, but our result shows that viral E6 is also a target of miR-34a in HPV infected cervical cancer cell lines.

It has been shown earlier that Cdt2 is an important cell cycle protein and is essentially required for hyperproliferation and survival of HPV positive cervical cancer cells, therefore destabilization of E6 by miR-34a reduces the level of Cdt2 which critically slows down growth and proliferation of HPV positive cervical cancer cells (Fig.3A, 3B and 4A 10x magnification). On the other hand, in the non-cancerous cell line (HEK293T) there is negligible suppression of Cdt2 upon overexpression of miR-34a, the proliferation rate catches up within 72 hours to normalcy (Fig. 3C). Change in cellular morphology could be seen in miR-34a overexpressed HPV positive cancerous cells (Fig.4A) indicating that Cdt2 suppression is causing cellular senescence probably due to halted G1 to S phase transition (Kiran *et al*, 2018) while non-cancerous cells behaved normal (Fig S2). This suppression in proliferation of HPV positive cervical cancer cell lines can be compensated by overexpression of either Cdt2 or E6 (Fig. 3D), indicating inverse relation between the miR-34a expression and cervical cancer cell proliferation. Additionally, miR-34a overexpression inhibits the cell invasion and migration abilities of cervical cancer cells (Fig.5A-5F) by inhibiting cell proliferation and inducing cell senescence (Bommer *et al*, 2007; He *et al*, 2007; Raver-Shapira *et al*, 2007; Tarasov *et al*, 2007; Tazawa *et al*, 2007).

E6 is an oncogenic protein that either suppresses or leads to degradation of several tumor suppressor proteins and checkpoint systems, which in turn leads to transformation and hyperproliferation of HPV infected cells. It also promotes stability of oncogenic cell cycle factor like Cdt2, empowering re-replication. miR-34a which happens to be a tumor suppressor miRNA is downregulated in many cancers including cervical cancer. Our study is the first to show that an anti-tumorous microRNA, miR-34a suppresses viral E6 protein and destabilizes overexpressed oncoprotein Cdt2 in HPV positive cell lines and stops their proliferation eventually by killing the HPV positive cancerous cells while no effect is seen on HPV negative cells. Since 99% cervical cancers are due to high-risk HPV infection (Moody & Laimins, 2010), our study opens up the potential of using miR-34a for HPV infected cervical cancer therapy which could be much cheaper, noninvasive and specific treatment plan for such fatal cancer.

## Acknowledgement

The authors are thankful to the Director Prof. A.K. Tripathi and Coordinator Prof. S.M. Singh, School of Biotechnology, Institute of Science, Banaras Hindu University for providing space and facilities to conduct the research. We thank the Central Discovery Center (CDC) for facilitating the super-resolution microscopy, maintained by SATHI BHU. We are thankful to Department of Biotechnology, Govt. of India for funding Samarendra K Singh (SKS), University Grants Commission (UGC), Govt. of India for funding Garima Singh and the Council of Scientific and Industrial Research (CSIR), New Delhi for funding Sonika Kumari Sharma. We also thank Dr. Arpita Singh, IMS, BHU for suggestions and critical review of the research.

## Authors’ Contribution

Samarendra K Singh was involved in the conceptualization and designing; supervision; writing - critical review & editing of the manuscript. Garima Singh was involved in investigation; methodology; project administration; data analysis; writing - original draft; review & editing of the manuscript. Sonika Kumari Sharma contributed to the methodology and writing--original draft & editing of the manuscript.

## Competing Interest

The authors declare no competing interest.

## Funding Information

The research was funded by Department of Biotechnology (DBT), Govt. of India, RLS grant (BT/RLF/Re-entry/43/2016) and Institute of Eminence (IOE) seed grant from Banaras Hindu University (BHU) to Samarendra K Singh. The University Grants Commission (UGC), Govt. of India also supported this research by funding Garima Singh and the Council of Scientific and Industrial Research (CSIR) by funding Sonika Kumari Sharma.

## Notes

### Competing Interest Statement

The authors have declared no competing interest.

