## Supplementary figures and images for "miR-34a negatively regulates cell cycle factor Cdt2/DTL in HPV infected Cervical Cancer Cells"

### Supplementary Figure

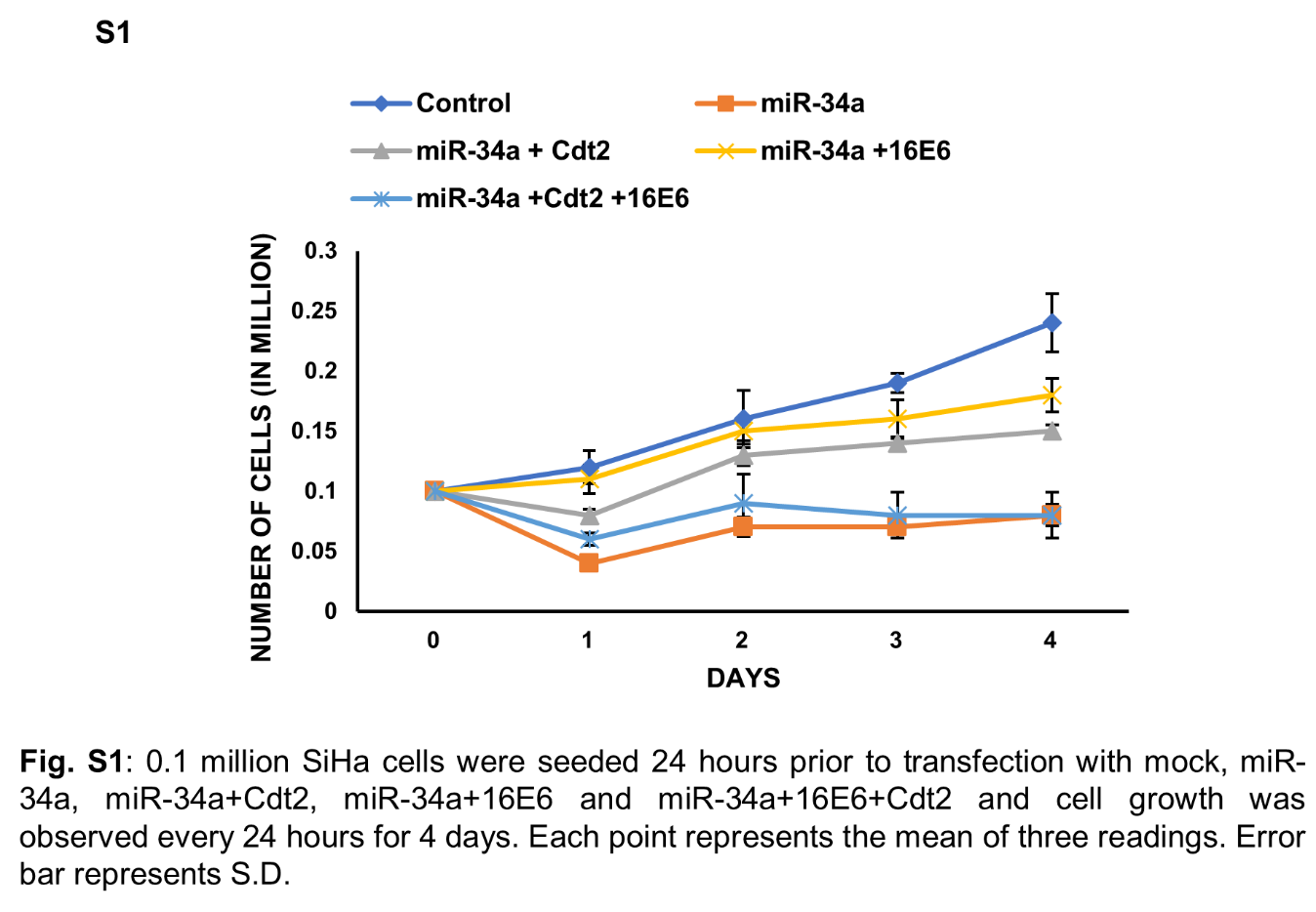

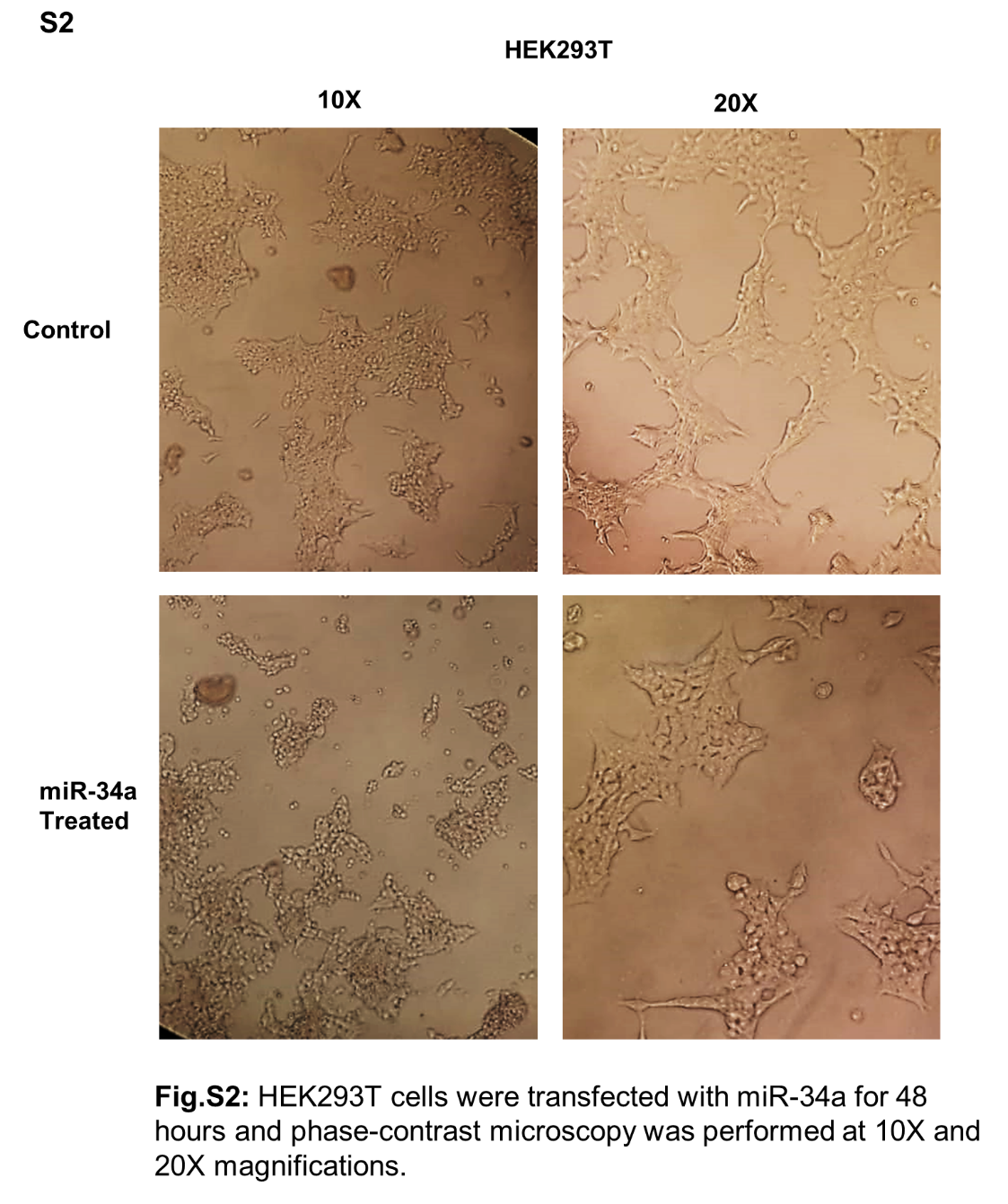
